# Homogenization of species composition and species association networks are decoupled

**DOI:** 10.1101/265264

**Authors:** Daijiang Li, Timothée Poisot, Donald Waller, Benjamin Baiser

**Keywords:** beta diversity, long-term changes, species co-occurrence, species composition, species interactions, networks

## Abstract

Ecological communities are comprised of both species and the biotic relationships among them. Biotic homogenization in species composition (i.e. increased site-to-site similarity) is recognized a common consequence of global change, but less is known about how species relationships change over space and time. Does homogenization of species composition lead to homogenization of species relationships or are the dynamics of species relationships decoupled from changes in species composition? To answer this question, we used long-term resurvey data to analyze changes in plant species association patterns between the 1950s and 2000s at 266 sites distributed among three community types in Wisconsin, USA. We used species associations (quantified via local co-occurrence patterns) as a proxy for species relationships. Species pairs that co-occur more/less than expected by chance have positive/negative associations. Shifts in species associations consistently exceeded the shifts observed in species composition. Less disturbed forests of northern Wisconsin have converged somewhat in species composition but not much in species associations. In contrast, forests in central Wisconsin succeeding from pine barrens to closed-canopy forests have strongly homogenized in both species composition and species associations. More fragmented forests in southern Wisconsin also tended to converge in species composition and in the species negative associations, but their positive associations diverged over the last half century. We conclude that long-term shifts in species relationships may be decoupled from those of species composition despite being affected by similar environmental variables.

## Introduction

Global environmental changes including shifts in climate, land use and management, and species invasions are affecting many communities and ecosystems (Vitousek *et al*., 1997; Sala *et al*., 2000) and forming novel ecosystems (Hobbs *et al*., 2009). Global biotic homogenization (BH) is occurring as sites converge in their species composition (McKinney & Lockwood, 1999; Olden & Poff, 2003). This has been documented in several ecosystems, taxonomic groups, and spatial scales (e.g., Rooney *et al*., 2004; Baiser *et al*., 2012; Li & Waller, 2015; Solar *et al*., 2015). Such declines in beta diversity adversely affect ecosystem functions (Olden *et al*., 2004) by reducing ecosystem services “insurance” effects (Loreau *et al*., 2003).

Environmental changes can also modify relationships among species (e.g. interactions, Tylianakis *et al*., 2008; Blois *et al*., 2013). This may result in new predator-prey interactions (Rockwell *et al*., 2011), intensified predation (Harley, 2011), changes in plant phenology leading to pollination mismatches (Hegland *et al*., 2009), and changes in non-trophic relationships among species such as species spatial association (Milazzo *et al*., 2013; Li & Waller, 2016). Species relationships may in fact be more sensitive and susceptible to environmental change, allowing them to act as a better indicator of change than species richness or composition (Tylianakis *et al*., 2008; Poisot *et al*., 2017). For example, relationships between a host and its parasites in the tropics changed in response to habitat modification without changes in species composition (Tylianakis *et al*., 2007). Species relationships also play critical roles in maintaining biodiversity and ecosystem functions at both local and regional scales (Bascompte *et al*., 2006; Gotelli *et al*., 2010; Harvey *et al*., 2017). As a result, monitoring species composition *and* species relationships simultaneously may provide a better understanding of how global change affects ecosystem structure and function (McCann, 2007; Valiente-Banuet *et al*., 2015).

Recently, there has been an upsurge of interest in species relationships in the context of ecological networks (McCann, 2007; Morales-Castilla *et al*., 2015; Tylianakis & Morris, 2017). This reflects important advances in the theory and methods of network analysis as well as its clear applications to conservation biology and restoration ecology (Cumming *et al*., 2010; Tylianakis & Morris, 2017). Ecological networks are composed of nodes and links where species are nodes and the relationships between them are links. Ecological networks provide a useful conceptual framework for studying species relationships and the complexity of biological systems. Nevertheless, most previous studies of species relationships focus on spatial variation in network structures, typically along some environmental gradient, rather than how these change over time (e.g. Mokross *et al*., 2014). Without long-term baseline data, it is difficult to study how species relationship networks may vary over time (Laliberté & Tylianakis, 2010; Poisot *et al*., 2015). However, given the rapid change in abiotic and biotic conditions across ecosystems worldwide (Tylianakis *et al*., 2008), exploring the temporal dynamics of species relationships is necessary for assessing biodiversity under global change.

Plant-plant relationships (e.g. facilitation, competition) form the foundation of plant community assembly, on which other types of relationships (e.g. trophic interactions in food webs, pollination interactions, host-parasite) build. Although plant-plant relationships are fundamental, they have received less attention than other types of ecological relationships. Part of the reason is that, while most other types of relationships can be detected by observations (e.g. pollination, predation, parasitism), plant-plant relationships are difficult to observe and thus require experiments to quantify. Conducting such experiments quickly becomes intractable as the number of possible relationships scales with the square of the number of species.

As an alternative to doing factorial experiments to detect how plant species interact, we can also examine how species associate spatially. Species association patterns are regularly used in community ecology and biogeography as a proxy for species relationships (Gotelli, 2000). Previous studies have suggested that biotic interactions are the main driver of local species associations (Morales-Castilla *et al*., 2015). Species association at the local scale is possible *despite or because of* interactions with other species in the community after passing the environmental filters. Consequently, we should be able to infer relationships among species based on how they co-occur locally (Gotelli, 2000; Araújo *et al*., 2011; Harris, 2016). Here, we use local-scale species association networks as proxies to represent species relationship networks. We use both terms interchangeably while acknowledge that we are not studying these relationships *directly*.

To study long-term changes in plant-plant species relationships, we applied network analysis to three forest plant community types in Wisconsin, USA, sampled first in the 1950s then again in the 2000s. Biotic homogenization in species composition has been observed in almost all of the plant communities in Wisconsin (Rooney *et al*., 2004; Rogers *et al*., 2008; Li & Waller, 2015). To understand whether plant-plant species relationships underwent similar homogenization in these communities, we examine long-term shifts in their species relationship networks here as inferred from species association patterns. We also ask whether species composition and relationships in these communities are affected by the same environmental variables in both time periods. Species associations may be inherently more labile than species composition (Poisot *et al*., 2017) reflecting the large number of interactions present among species (with *n* species, we have (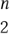) possible species associations). Therefore, we hypothesize that changes in species relationship networks are decoupled from changes in species composition. Specifically, we do not expect biotic homogenization in species composition to necessarily be reflected in changes in species association networks. We also hypothesize that species composition and relationships are driven by the same environmental factors given the fact that relationships are build on species identities. However, we do not expect these environmental factors to have the same effect over time given observed changes in land use and climate across these communities. In sum, we sought to demonstrate whether species relationships and species composition have similar responses to global environmental change.

## Methods

### Vegetation data

In the 1950s, John Curtis and his students and colleagues canvassed the state of Wisconsin to find the best remaining examples of natural vegetation then sampled 1000+ sites and diverse community types (Curtis, 1959). They only chose sites with no obvious disturbances and their plots were always at least 30m away from any edges. Within each site, they recorded the presence and absence of all vascular plants in each of many sampled 1-*m*^2^ quadrats. The number of quadrats sampled at each site varied but was usually 20. They were careful to archive all their original data in the Plant Ecology Laboratory at University of Wisconsin – Madison (http://www.botany.wisc.edu/PEL/, Waller *et al*., 2012). Here, we use data from three community types re-surveyed since 2000 using similar methods: northern upland forests (NUF, 108 sites), central sands pine barrens forests (CSP, 30 sites), and southern upland forests (SUF, 128 sites). Because the original sites were not permanently marked, we analyze data from these “semi-permanent” plots. The resurveys sampled 2-6 times as many quadrats per site as the original survey. All taxonomy was carefully synchronized between periods. To allow fair comparisons with matched sampling effort, we randomly sub-sampled the the 2000s survey data using the same number of quadrats as used in the 1950s. Collectively, we analyzed species presence/absence data from >5000 quadrats distributed among the same 266 sites in the two time periods.

### Environmental data

We analyzed a suite of environmental variables to detect drivers of plant composition and association patterns. These variables fall into several major categories including: soil properties, landscape variables, canopy shade, and climatic variables. We used Principal Component Analysis (PCA) to reduce the dimensionality of the soil variables (including soil nitrogen, proportions of sand and clay, pH, etc.) yielding two primary axes (`soil_pc1, soil_pc2`). We also used PCA to compute 2-axis summaries of landscape variables (`landsc_pc1, landsc_pc2`) for all sites except those in the Central Sand Plains. These variables were: proportions of different land use types, road density, and house density, all computed for the area within 2km of each site. We did not have shade data for the northern and southern upland forests. We extracted local climatic variables based on average daily precipitation and minimal temperature for all sites using a Wisconsin climate database that covers 1950 to 2006 (Kucharik *et al*., 2010). These data derive from an extensive network of weather stations distributed throughout the state. Downscaled data were generated via spatial interpolation. To represent each period of sampling, we averaged climate variables over two five year periods: 1950-1954 and 2002-2006. This accounts for potential lags in species’ responses and inter-annual climatic variation.

### Plant association networks

Within each vegetation type and time period, we constructed a quadrat by species matrix with rows for each quadrat (nested within a site) and columns for each species. Values in the cells of this matrix reflect the presence or absence (1/0) of that species in that quadrat. We treat the 1-*m*^2^ quadrat as the sample unit here because plants that co-occur at this scale are most likely to also interact. We removed species occupying fewer than six quadrats at each period to exclude rare species and facilitate the determination of core species co-occurrence pairs. We then used this quadrat by species matrix to infer species pairs that are more or less likely to co-occur with each other as compared to random expectation according to two methods.

Our first method is based on the traditional null model approach (Gotelli, 2000), which is commonly used to study species co-occurrence patterns. We calculated the partial C-score for each pair of species as (*c_i_* - *m_ij_*)(*c_j_* - *m_ij_*), where *c_i_* and *c_j_* are the number of quadrat occurrences of species *i* and *j* and *m_ij_* is the number of quadrats where both species occurred. We then shuffled the cells of the quadrats by species matrix 5000 times using the fixed-fixed randomization algorithm. This null model maintains row and col sums (species richness within each quadrat and species frequency across all quadrats) of the matrix. In each iteration, partial C-scores for all species pairs were computed, generating a null distribution from the 5000 randomizations. This was then used to judge whether the observed C-score reflects higher or lower co-occurrence than expected by chance. For more details, see Li & Waller (2016).

A recent study concluded that this null model approach has relatively low power to infer true species interactions from co-occurrence patterns, suggesting the use of Markov networks instead (Harris, 2016). Unfortunately, current implementations of Markov networks are restricted to 20 species or fewer (Harris, 2016) thus we could not apply them to our dataset. Instead, we use generalized linear models (GLMs) as our second method. GLMs have similar power as Markov networks (Harris, 2016) but extend to include many more species. For the GLMs, we fitted Bayesian regularized logistic regression to the presence/absence for each species (response) using the presence/absence of other species as predictors. This method generates two regression coefficients and p-values for each species pair. We averaged these to estimate the strength of species interactions (cf. Harris, 2016). The two methods we employed yielded qualitatively similar results that both support our main conclusions. We therefore only report results from GLMs in the main text as these may have higher statistical power. For the C-score null model results, see the Appendix.

With the list of positive and negative association species pairs, we built one positive association metaweb and one negative association metaweb for each vegetation type and time period. We then built positive and negative association networks for each site from these two metawebs by sub-setting the species observed at that site. This assumes that species relationships between species do not differ across sites of the same vegetation type and time period. It therefore results in more conservative results. To remove this assumption, one can also build an association network for each site independently. However, we lacked the power to do this given the limited number of quadrats (mostly 20) per site. Therefore, the association networks for each site in our analyses were derived from the metawebs instead of build independently.

## Data analysis

### Changes in spatial *β* diversity over time

We first calculated pairwise beta diversity in both species composition and species association networks within each vegetation type and time period using the methods proposed by Legendre & De Cáceres (2013). For pairwise beta diversity of species composition, the input is a site by species matrix, with species abundances in the cells; for species association pairwise beta diversity, the input is a site by species pairs (non-random pairs inferred via methods described above) matrix. In this way, we treat each non-random species pair as a “species” in traditional community ecology analyses (cf. Poisot *et al*., 2017). We have also calculated pairwise beta diversity in species association networks using the method proposed by Poisot et *al*. (2012). This method divides beta diversity of interaction networks (*β_WT_*) additively into *β_ST_* (dissimilarity of interactions due to species turnover) and *β_OS_* (dissimilarity of interactions established between species common to both networks). As we are forced to assume that species relationships do not change across sites within each vegetation type and time period (i.e., *β_OS_* = 0), the calculated *β_WT_*, a measure of beta diversity for site pairs, correlated tightly with those calculated from methods proposed by Legendre & De Cáceres (2013) (Pearson *r* > 0.98 for NUF sites as an example). Thus, we only use the methods proposed by Legendre & De Cáceres (2013) to calculate pairwise beta diversity. This approach allows us to directly compare pairwise beta diversity for species composition to beta diversity for species associations as both are calculated the same way. We compared pairwise beta diversity of each vegetation type between the 1950s and the 2000s using Wilcox paired tests. Lower (or higher) beta diversity in the 2000s suggests biotic homogenization (or differentiation).

Given the number of sites in NUF (108) and SUF (126), we have a large number of pairwise beta diversity measures (5778 and 7875, respectively). Such large sample sizes make it possible to obtain statistically significant results that may not be biologically significant. Therefore, we also test changes in beta diversity for each vegetation type using a distance-based permutational test for homogeneity of multivariate dispersion (PERMDISP, Anderson *et al*., 2006). PERMDISP calculates distance of each site to the centroid of the ordination space and then tests whether these distances are different across groups (i.e. 1950s vs 2000s) with permutation tests. We also use results from PERMDISP to visualize species composition and association patterns for each vegetation type and time period.

### Within site changes over time

To compare rates of change in species composition and species relationships over time, we calculated beta diversities between periods within each site (i.e. a site in the 1950s vs the same site in the 2000s), again using the Legendre & De Cáceres (2013) method. We then used paired t-test to examine whether these beta diversities reflecting changes in species associations significantly exceeded the beta diversities reflecting changes in species composition (i.e., whether species associations changed more than species composition). To confirm these results, we also applied Permutational Multivariate Analysis of Variance (PERMANOVA) to compare changes in species composition and species associations.

### Environmental drivers

To understand environmental drivers for species composition and association networks, we conducted distance-based redundancy analysis (RDA) for each vegetation type and time period. We transformed species composition and association matrix into distance matrices with the Hellinger index (Legendre & Legendre, 2012). We used environmental variables as predictors in RDAs. To identify the most significant environmental variables, we used forward variable selection and AIC-based statistics over 999 replicate runs for each matrix (c.f. Poisot et *al*., 2017). The order of variable selection provides insight into the importance of environmental variables with the earlier selected variables generally affecting species composition or the associations more. This analysis allowed us to study whether species composition and association are affected by similar sets of environmental variables in the same way and how these relationships changed over time. All analyses were conducted in R v3.4.0 (R Core Team, 2017).

## Results

Across all vegetation types, the majority (>90%) species pairs co-occurred randomly at both time periods as inferred from the results using Bayes GLMs (Table 1). Among the non-random species pairs, more species pairs co-occurred positively than negatively across all vegetation types and time periods (Table 1).

**Table 1:** Summary of species used and species associations for each vegetation type and time period. Abbreviations: NUF, northern upland forests; CSP, central sand plains; SUF, southern upland forests.

| Type | Date | Species Used | Pairs (Neg. Assoc.) | Pairs (Pos. Assoc.) | Pairs (Random) | Total Pairs |
| --- | --- | --- | --- | --- | --- | --- |
| NUF | 1950s | 146 | 91 (0.9%) | 393 (3.7%) | 10101 (95.4%) | 10585 |
| NUF | 2000s | 160 | 140 (1.1%) | 473 (3.7%) | 12107 (95.2%) | 12720 |
| CSP | 1950s | 61 | 31 (1.7%) | 83 (4.5%) | 1716 (93.8%) | 1830 |
| CSP | 2000s | 55 | 19 (1.3%) | 72 (4.8%) | 1394 (93.9%) | 1485 |
| SUF | 1950s | 225 | 220 (0.9%) | 771 (3.1%) | 24209 (96.1%) | 25200 |
| SUF | 2000s | 186 | 124 (0.7%) | 522 (3%) | 16559 (96.2%) | 17205 |

In the NUF region, species composition and the positive species associations had similar levels of dispersion in both time periods within the ordination space (Fig. 1), suggesting that homogenization did not occur. This was confirmed by PERMDISP results (permutation test, all *p* > 0.25, Table 2). However, pairwise site beta diversity calculated with methods proposed by Legendre & De Cáceres (2013) suggested no changes in positive associations but homogenization in species composition and differentiation in negative associations (Paired Wilcoxon test, both *p* < 0.001, Table 2).

**Figure 1:**
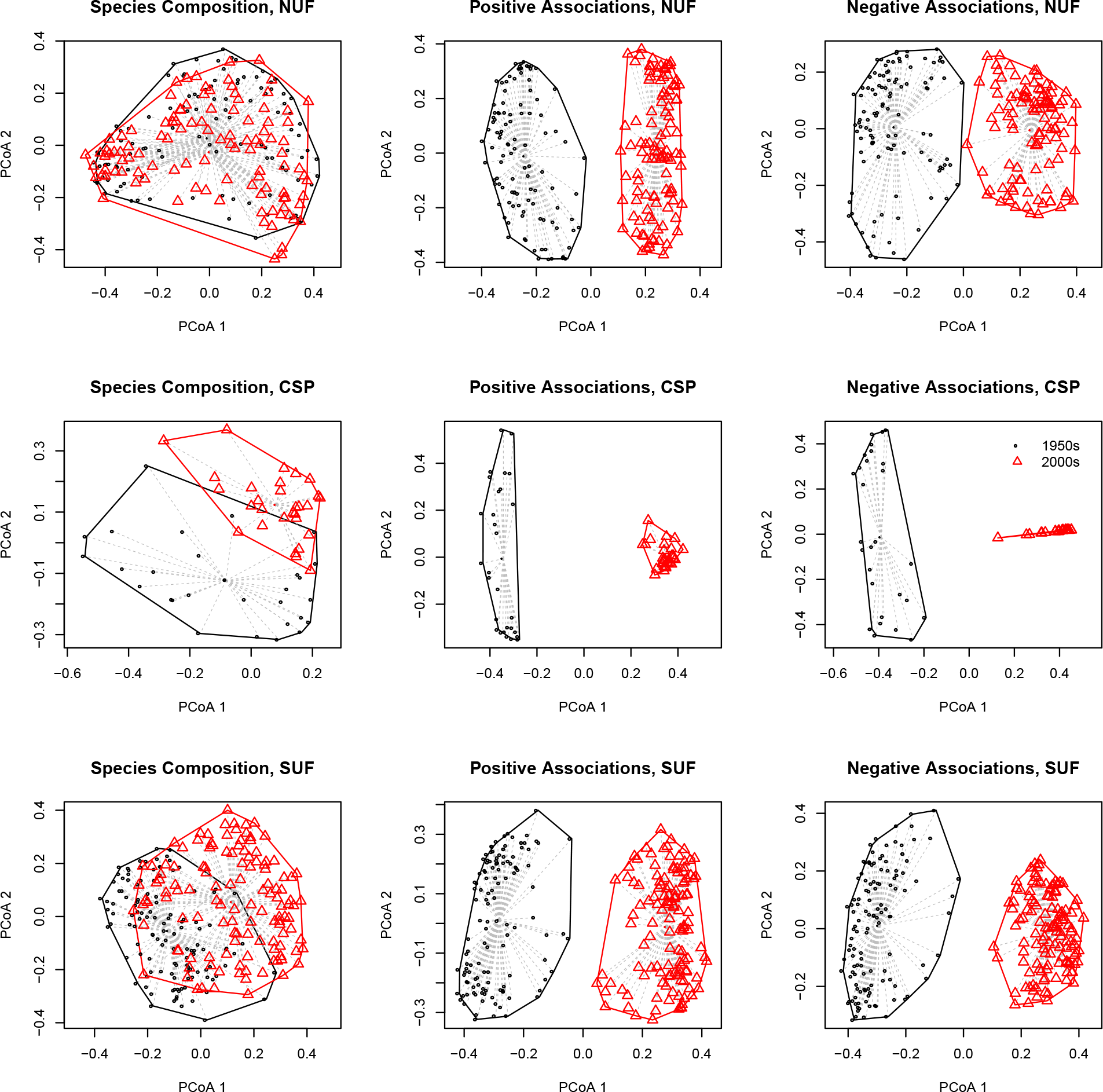
Ordination of species composition and associations. Circles represent sites in the 1950s while triangles represent sites in the 2000s. Note that although the periods overlapped considerably in species composition, significant species associations in the two time periods did not overlap at all in any vegetation type. This suggests that species relationships differ between the two time periods in all vegetation types. The spread of sites within each time period / vegetation type reflects variation in species relationships across sites.

**Table 2:** Homogenization in species composition does not always result in the homogenization of species association networks. Numbers in the ave_1950 (pairwise) and ave_2000 (pairwise) columns represent average pairwise beta diversity between sites within the same vegetation type in the 1950s and the 2000s, respectively. Numbers in the ave_1950 (permdisp) and ave_2000 (permdisp) columns represent average distance to group centroid in the ordinations of sites within the same vegetation type in the 1950s and the 2000s, respectively. The column direction (pairwise) indicates directions of changes in the pairwise site beta diversity over time, tested with paired Wilcoxon tests. The column direction (permdisp) indicates directions of changes in the distance between sites and the ordination centroid over time, tested with permutation tests. Abbreviations: Spe. Comp, species composition; Pos. Assoc., positive associations; Neg Assoc., negative associations, homog., homogenization; diff., differentiation, *, p < 0.05; **, p < 0.01; ***, p < 0.001.

| vegtype | var2 | ave_1950<br>(pair-<br>wise) | ave_2000<br>(pair-<br>wise) | ave_1950<br>(per-<br>mdisp) | ave_2000<br>(per-<br>mdisp) | direction<br>(pair-<br>wise) | direction<br>(per-<br>mdisp) |
| --- | --- | --- | --- | --- | --- | --- | --- |
| NUF | Spe.<br>Comp. | 0.595 | 0.588 | 0.513 | 0.509 | homog.*** | no<br>change |
| NUF | Pos.<br>Assoc. | 0.755 | 0.757 | 0.545 | 0.547 | no<br>change | no<br>change |
| NUF | Neg.<br>Assoc. | 0.766 | 0.787 | 0.554 | 0.568 | diff.*** | no<br>change |
| CSP | Spe.<br>Comp. | 0.447 | 0.299 | 0.418 | 0.327 | homog.*** | homog.*** |
| CSP | Pos.<br>Assoc. | 0.643 | 0.398 | 0.463 | 0.285 | homog.*** | homog.*** |
| CSP | Neg.<br>Assoc. | 0.699 | 0.443 | 0.513 | 0.323 | homog.*** | homog.*** |
| SUF | Spe.<br>Comp. | 0.572 | 0.546 | 0.516 | 0.497 | homog.*** | homog.* |
| SUF | Pos.<br>Assoc. | 0.712 | 0.744 | 0.511 | 0.538 | diff.*** | diff.** |
| SUF | Neg.<br>Assoc. | 0.769 | 0.713 | 0.553 | 0.517 | homog.*** | homog.*** |

In the CSP region, sites converged in ordination space in the 2000s when compared to 1950s (Fig. 1). This suggests that these sites experienced homogenization in both species composition and species associations (positive and negative). Results from both Paired Wilcoxon test on pairwise beta diversity and PERMDISP (all *p* < 0.001, Table 2) confirm this interpretation.

Sites in the SUF region also showed a tendency to converge in species composition and negative associations between periods (Fig. 1). In contrast, positive associations tended to diverge. These results were supported by the parallel analyses of pairwise site beta diversity and PERMDISP (all *p* < 0.001 for paired Wilcoxon tests and all *p* < 0.035 for PERMDISP, Table 2).

For all regions, shifts in beta diversity for the species association networks exceeded those for species composition (*p* = 0.001 within each site Fig. 2). Thus, it appears that species association networks have changed faster than species composition. Large turnover in species associations (positive or negative) may reflect only slight changes in species composition.

**Figure 2:**
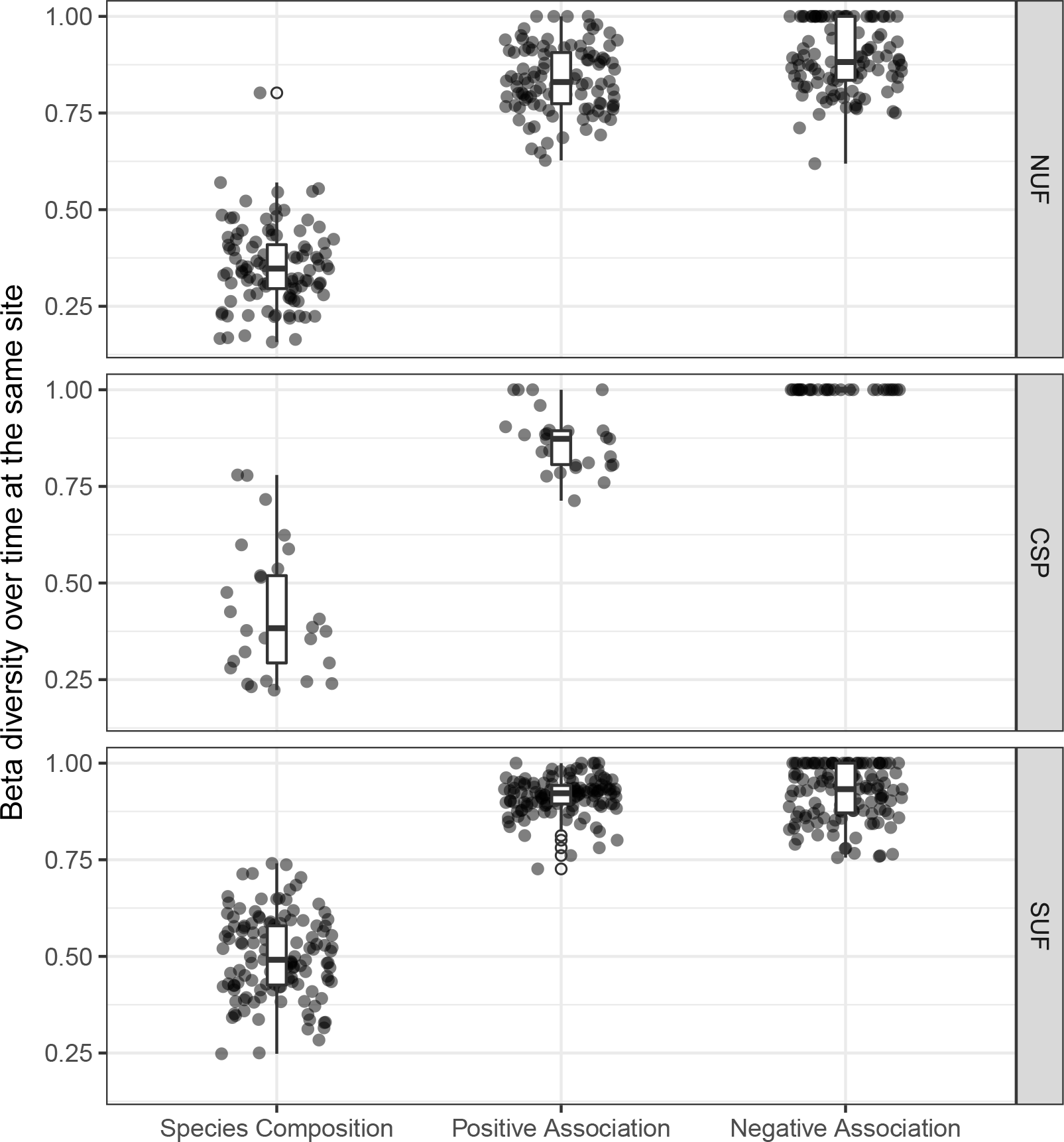
Changes in species composition and association of the same site over time. Each point represents changes in beta diversity at the same site between the 1950s and the 2000s. Turnover in species relationships appear to be much greater than the turnover in species composition.

Species composition and species associations were largely influenced by the same set of environmental variables within each vegetation type and each time period (Table 3). For CSP and SUF in the 1950s, species composition and species association networks were affected by almost the same set of environmental variables. This pattern still holds for the SUF in the 2000s but less so for the CSP and the NUF sites. More importantly, the importance of environmental variables on plant communities has changed over time. For example, shade was most important for the CSP sites in the 1950s but lost importance by the 2000s. Soil properties there strongly affected the SUF sites in the 1950s became less important by the 2000s while climatic variables gained importance. Precipitation had the second most important effect on species composition of NUF in the 1950s but became least important in the 2000s.

**Table 3:** Selected environmental variables for species composition and species associations. Numbers are the order that variables are selected. ‘-’ means that the environmental variable is not available. Blank cells mean that these variables are dropped out.

| type | date | var | shade | precip | tmin | soil_pc1 | soil_pc2 | landsc_pc1 | landsc_pc2 |
| --- | --- | --- | --- | --- | --- | --- | --- | --- | --- |
| NUF | 1950s | Spe. Comp. | - | 2 | 4 | 1 | 3 |  | 5 |
|  |  | Pos. Assoc. | - | 4 | 2 | 1 | 3 |  |  |
|  |  | Neg. Assoc. | - |  | 1 | 2 | 3 |  | 4 |
|  | 2000s | Spe. Comp. | - | 5 | 1 | 2 | 3 |  | 4 |
|  |  | Pos. Assoc. | - | 4 | 3 | 1 | 2 |  | 5 |
|  |  | Neg. Assoc. | - | 5 | 1 | 3 | 4 |  | 2 |
| CSP | 1950s | Spe. Comp. | 1 |  |  |  | 2 | - | - |
|  |  | Pos. Assoc. | 1 |  |  |  | 2 | - | - |
|  |  | Neg. Assoc. | 1 |  |  |  | 2 | - | - |
|  | 2000s | Spe. Comp. |  | 4 | 2 | 3 | 1 | - | - |
|  |  | Pos. Assoc. |  | 1 |  | 3 | 2 | - | - |
|  |  | Neg. Assoc. |  | 1 |  |  | 2 | - | - |
| SUF | 1950s | Spe. Comp. | - | 4 | 3 | 2 | 1 | 5 |  |
|  |  | Pos. Assoc. | - | 4 | 3 | 2 | 1 | 5 |  |
|  |  | Neg. Assoc. | - | 5 | 3 | 2 | 1 |  | 4 |
|  | 2000s | Spe. Comp. | - | 1 | 2 | 3 | 4 | 5 |  |
|  |  | Pos. Assoc. | - | 2 | 1 | 3 | 4 | 5 |  |
|  |  | Neg. Assoc. | - | 1 | 2 | 3 | 4 |  |  |

## Discussion

Few studies have quantified changes in both species composition and species relationship networks over time (Burkle *et al*., 2016). It takes considerable effort to construct a single interaction network, let alone networks at multiple sites over two or more time periods. We found only two empirical studies on homogenization of ecological networks. Laliberté & Tylianakis (2010) found that deforestation homogenized parasitoid-host networks in tropical areas. Although they have temporal data of parasitoid-host networks (monthly samples for 17 months), their main conclusion was derived from spatial comparisons among different land use categories. Kehinde & Samways (2014) examined biotic homogenization of insect-flower interactions in vineyards managed under agri-environmental schemes in the Cape Floristic Region. They found no evidence of homogenization for interaction networks when comparing vineyards to natural sites.

Our study may thus be the first to explore temporal changes in beta-diversity of species relationship networks. By using patterns of species co-occurrence to indicate species relationships and by using a valuable, high-resolution, long-term dataset, we were able to examine parallel changes in both community composition and species interaction networks. We found species relationship networks can homogenize, differentiate, or show no change through time in different vegetation types regardless of the homogenization dynamics of species composition. Long-term changes in species composition and species interactions thus appear to be decoupled.

In the northern upland forests (NUF) of Wisconsin, plant communities were relatively stable in terms of their species composition and species associations. Compared to other community types, NUF area has a lower human population, less land use change, and less habitat fragmentation. In this study, we found no overall changes in beta diversity of species composition and species associations when tested with PERMDISP. However, paired Wilcoxon test on beta diversity between site pairs suggested significant homogenization in species composition, matching the conclusion (biotic impoverishment and homogenization) reached in a previous study of a subset of these sites (Rooney et *al*., 2004). Here, we also found no changes in the among-site diversity of positive species associations but significant differentiation in negative species associations. Given the large number of sites in NUF (108), the results of Wilcoxon tests may not be biologically significant despite their statistical significances. These results suggest that shifts in species composition may not occur in the same direction as shifts in species relationships.

Plant communities in the central sand plains (CSP) historically were fire-maintained pine barrens with open canopies. However, they are succeeding into close-canopy upland forests because of fire suppression (Li & Waller, 2015). Fire suppression has resulted in homogenization in both species composition (Li & Waller, 2015) and functional trait composition (Li & Waller, 2017). This probably reflects declines in habitat heterogeneity within these communities. In the 1950s, sites in the CSP had different canopy coverage, forming mosaics of burned and unburned habitats to support different plant communities. In the 2000s, however, sites were similar with each other in their canopy cover due to fire suppression and succession, filtering out shade intolerant species (Li & Waller, 2017) and homogenizing plant communities (Li & Waller, 2015). Given these ecological changes, it is not surprising to find significant homogenization in both positive and negative species associations in these communities.

Sites in the southern upland forests (SUF) have been affected by development and land use changes more than any other plant community in Wisconsin (Rogers *et al*., 2008). Currently, most of these sites are fragmented and disturbed by nearby anthropogenic activities including roads, development, and agriculture. Previous studies suggest that habitat degradation and fragmentation tend to homogenize species composition by decreasing species diversity, which can also reduce network complexity and stability (Tylianakis *et al*., 2007; Laliberté & Tylianakis, 2010; Mokross *et al*., 2014). However, the fact that habitat fragmentation can result in greater differences in interaction network structure (Bordes *et al*., 2015) suggests that network simplification (less nodes and/or edges) does not necessarily cause network homogenization. Networks can differ across sites if individual networks contain unique interactions even if they show a general trend toward simplification. In these SUF communities, the importance of species dispersal limitation and stochastic factors have increased while the importance of species interactions have decreased over time (Li & Waller, 2016). It is thus likely that stochastic assembly processes are forming novel sets of interaction among species even though species diversity has decreased. Indeed, we found on average 116 significant positive species pairs per site in the 1950s, but only 63 significant positive pairs per site in the 2000s. Therefore, association network at each site was simpler in the 2000s (paired t-test, *df* = 125, *t* = 11.90, *p* < 0.0001). However, both Wilcoxon paired test on pairwise beta diversity of positive association networks and PERMDISP suggested that positive association networks in the 2000s have differentiated since the 1950s (Table 2). Therefore, in the SUF, we found homogenization of species composition but differentiation of species positive associations, despite the fact that we did not find landscape variables to be the most important ones for interactions in the SUF (results from the null model did pick up landscape variables as the most important variable for positive species interactions, appendix table S3).

Although changes in species relationships are occurring faster than changes in species composition and appear decoupled from them, they appear to be affected by a similar set of environmental variables within each vegetation type and time period (Table 3). For example, species relationships and composition at the CSP in the 1950s were affected by the same set of environmental variables (canopy shade and soil); similar pattern found at the SUF and NUF even though with slightly different orders for environmental variables (cf. Poisot *et al*., 2017). However, the importances of environmental variables for species relationships and compositions have changed over time for all vegetation types, especially CSP and SUF. For example, canopy shade was the most important variables for CSP sites in the 1950s, but it was dropped out in the 2000s. This is understandable because sites in the CSP are close-canopy forests now with very little variation in canopy cover. In the SUF, soil properties used to the most important variables for both interactions and compositions, but they were replaced by climatic variables in the 2000s. For the relatively stable sites in the NUF, we did not find such abrupt changes in the importances of environmental variables over time. These results suggest that species interactions and compositions are mostly affected by the similar set of environmental variables (though not identical). As global environmental changes accelerate, the relative importance of environmental variables may also change, resulting in novel arrays of species and consequent species interactions (Blois *et al*., 2013; Milazzo *et al*., 2013).

Here, we used species association (co-occurrence) patterns to represent species relationships as it was intractable to quantify interactions among hundreds of plant species. Species associations may give false positive (hypothesized links that does not exist in real system) relationships between species and may not detect all real interactions (Delalandre & Montesinos-Navarro, 2018; Freilich *et al*., 2018). However, our main goal is to study whole community dynamics rather than to pinpoint exact interactions between particular species pairs. For this purpose, species spatial association network is a necessary and useful proxy (Freilich *et al*., 2018) because it can provide valuable information regarding the net output of direct and indirect effects among multiple plant species (rather than exact pairwise interactions, Delalandre & Montesinos-Navarro, 2018). Furthermore, co-occurrence networks can also predict overall community responses to disturbance (Tulloch *et al*., 2018). To reduce the potential for bias, we studied species association patterns at a fine spatial scale (1m^2^) and used statistical methods (Bayes GLMs) that have relatively high power for detecting species interactions (Harris, 2016). Furthermore, we used the null model method and reported its result in the appendix. Because both methods provide quantitatively similar results and reach the same conclusion, our conclusion that changes in species relationships and species composition are decoupled is likely to be robust. Consequently, our results about species association networks may also hold for species interaction networks.

A necessary assumption in this study is that species relationships within each vegetation type and time period are consistent across sites. In reality, relationships between species can vary through space and time (Poisot *et al*., 2015). Because we lacked the data to analyze how species relationships may have differed over sites, we pooled data across sites to gain insights into the average nature of species relationships within each community. This makes our results conservative as accounting for variation in species interactions over sites could only show greater variation in network structure, strengthening our conclusion that temporal changes in species relationships and composition are decoupled.

Our results suggest that species relationships may not be experiencing a general trend toward homogenization as novel relationships may be forming in response to rapid global change. This opens important questions into how these changes in species relationships may affect community stability, ecosystem services, and species co-evolution. Studies of beta diversity in species relationships remain in their infancy (Burkle *et al*., 2016). Given the importance of species interactions and their potential unpredictable relationship with species composition, future empirical and theoretical research that investigates patterns, causes, and consequences of changes in beta diversity of interaction networks are needed.

## Conclusion

Previous studies of long-term change in communities have mostly focused on species composition, often documenting biotic homogenization. Relationships among species, however, may be just as important in that they, too, can affect ecosystem function with important implications for conservation and restoration. Remarkably little is know about how species relationship networks shift in response to global environmental changes. Our study here, based on remarkably detailed long-term data, makes clear that these changes can be complex. Species relationship networks homogenized, differentiated, or remained unchanged among the three vegetation types. In contrast, species composition converged in all three. Although long-term changes in species composition and species relationships were decoupled in these communities, they were affected by similar sets of environmental variables. Their relative importance, however, changed between the 1950s and 2000s particularly in the more disturbed communities. As environmental changes accelerate, we may see fewer but unique relationships among species. Taken together, these results highlight the need to study species relationships in addition to species composition as changes in relationships may not necessarily be predictable from changes in species composition.

